# PINK1 Deficiency Alters Muscle Stem Cell Fate Decision and Muscle Regenerative Capacity

**DOI:** 10.1101/2023.06.23.546123

**Authors:** George Cairns, Madhavee Thumiah-Mootoo, Mah Rukh Abassi, Jeremy Racine, Nikita Larionov, Alexandre Prola, Mireille Khacho, Yan Burelle

## Abstract

Maintenance of optimal mitochondrial function plays a crucial role in the regulation of muscle stem cell (MuSC) behavior, but the underlying maintenance mechanisms remain ill defined. In this study, we explored the importance of mitophagy, as a mitochondrial quality control regulator, in MuSCs and the role this process plays in maintaining optimal muscle regenerative capacity. Here we show that MuSCs exhibit dynamic alterations in mitophagy under different physiological myogenic states. In particular, quiescent MuSCs exhibit high levels of PINK1/Parkin-dependent mitophagy, which is rapidly decreased upon transition to an early activation state. Genetic disruption of this pathway using *Pink1* knockout mice reduced mitophagy in quiescent MuSCs, which was accompanied by increased mitochondrial ROS release and mitochondrial network fragmentation. These abnormalities led to hampered self-renewal of MuSCs which ultimately translated in a progressive loss of muscle regeneration following repetitive injury. However, proliferation and differentiation capacity were unaltered in the absence of PINK1, indicating that altered fate decisions is the main mechanism underlying impaired muscle regeneration. Impaired fate decisions in PINK1 deficient MuSCs could be restored by scavenging excess mitochondrial ROS. Together, these data shed new light on the regulation of mitophagy during MuSC state transitions and position the PINK1-dependent pathway as an important regulator of MuSC mitochondrial properties and fate decisions.

## 1 INTRODUCTION

Skeletal muscle displays a remarkable plasticity which allows them to regenerate effectively. A key factor underlying this capacity is the existence of Muscle Stem Cells (MuSCs), a population of myogenic stem cells located in a specialized niche at the fiber surface where they remain in a reversible “dormant” state known as quiescence^1–3^. Upon injury, MuSCs activate and re-enter the cell cycle. As cells proliferate, a majority commits to the myogenic lineage to generate and repair tissue, while a small number retain their quiescent properties to renew the stem cell pool ^1–3^. Multiple studies have shown that an optimal balance between commitment and self-renewal is crucial to maintain muscle regenerative capacity ^1–3^, although the mechanisms through which this is achieved are not well understood.

Mitochondria have emerged as key players in the regulation of stem cell behavior, including their decision to commit or self-renew ^4–7^. More specifically, quiescent stem cells including MuSCs were shown to maintain a small pool of mitochondria that display dependence on fatty acid oxidation^8,9^, reduced oxidative phosphorylation activity^10–12^ and low ROS emission^10,12^, which allows them to emit pro-quiescence signals while limiting oxidative damage. Conversely, several studies have shown that moving away from these metabolic properties provokes stem cell commitment and is required for normal progression toward terminal differentiation^5–7,12^. As a result there is currently a strong impetus for understanding the mechanisms involved in regulating mitochondrial qualities, and how this impacts MuSC regenerative abilities.

Studies on autophagy have recently unveiled the critical role of this bulk recycling process in the maintenance of stem cell homeostasis and function^4,7,13^. More specifically, autophagy was shown to be essential for preserving the global integrity of cellular constituents, including mitochondria^13,14^, and to support the increased bioenergetic demands of proliferation through provision of energy substrates^15^. However, what is less understood is the role of specific autophagy mechanisms such as mitophagy in regulating mitochondrial qualities and MuSC states. Selective removal of mitochondria through mitophagy can occur by means of PINK1/Parkin-mediated ubiquitination of mitochondrial proteins and subsequent recruitment of the autophagy machinery, or through mitophagy receptors that can directly bind LC3 on nascent autophagosomes^4,16,17^. The contribution of these various pathways in MuSCs currently remains unknown. In this study, we therefore explored the regulation of mitophagy under different MuSC states, and used a mouse model of PINK1 deficiency to assess the role of the PINK1/Parkin pathway. Our results reveal that mitophagy is dynamically regulated during transition from quiescence to activation. Moreover, we show that PINK1 is dispensable for normal differentiation, but plays a particular role in maintaining an optimal balance of MuSC fate decisions through a mechanism that implicates mitochondrial ROS.

## 2 RESULTS

### 2.1 Mitophagy is prominent in quiescent MuSCs and downregulated during activation

MuSCs normally adopt a reversible state of quiescence, but can rapidly activate and proliferate in response to a variety of signals in the surrounding environment. To gain insight on the role of mitophagy in MuSCs, the regulation of this process was first examined using four transcriptomics datasets comparing quiescent and *in vitro* or *in vivo* activated MuSCs. One dataset (GSE103163) compared MuSC that were *in situ* fixed prior to isolation, to preserve their native quiescent state, to unfixed MuSC that had initiated *in vitro* activation during the 3-5h isolation process^18^. The three other datasets (GSE 70736, 55490, and 47177) compared MuSCs purified from uninjured and injured muscles 36-72h post injection of cardiotoxin or BaCl_2_ ^14,19,20^. Depending on the method and timing of activation, 12 to 37 of the 46 genes associated with mitophagy in Gene Ontology (GO:0000423; 1901524;1903146;0000422) were found in the list of differentially enriched genes in these datasets, representing a significant enrichment in all cases (Fig. 1A and C). Closer investigation of genes involved in ubiquitin-dependent mitophagy revealed that the expression of *Pink1* and *Parkin* was systematically enriched in quiescent MuSCs and downregulated in activated cells (Fig. 1B and D). Several genes encoding mitophagy receptors (*Fundc1, Bnip3, Bnip3L, Bcl2l13, Ambra1*) were also differently expressed, but the pattern of changes was less consistent across datasets. Together these results suggested that mitophagy pathways undergo rapid transcriptional changes in MuSCs transitioning from a quiescent to activated state.

**Figure 1:**
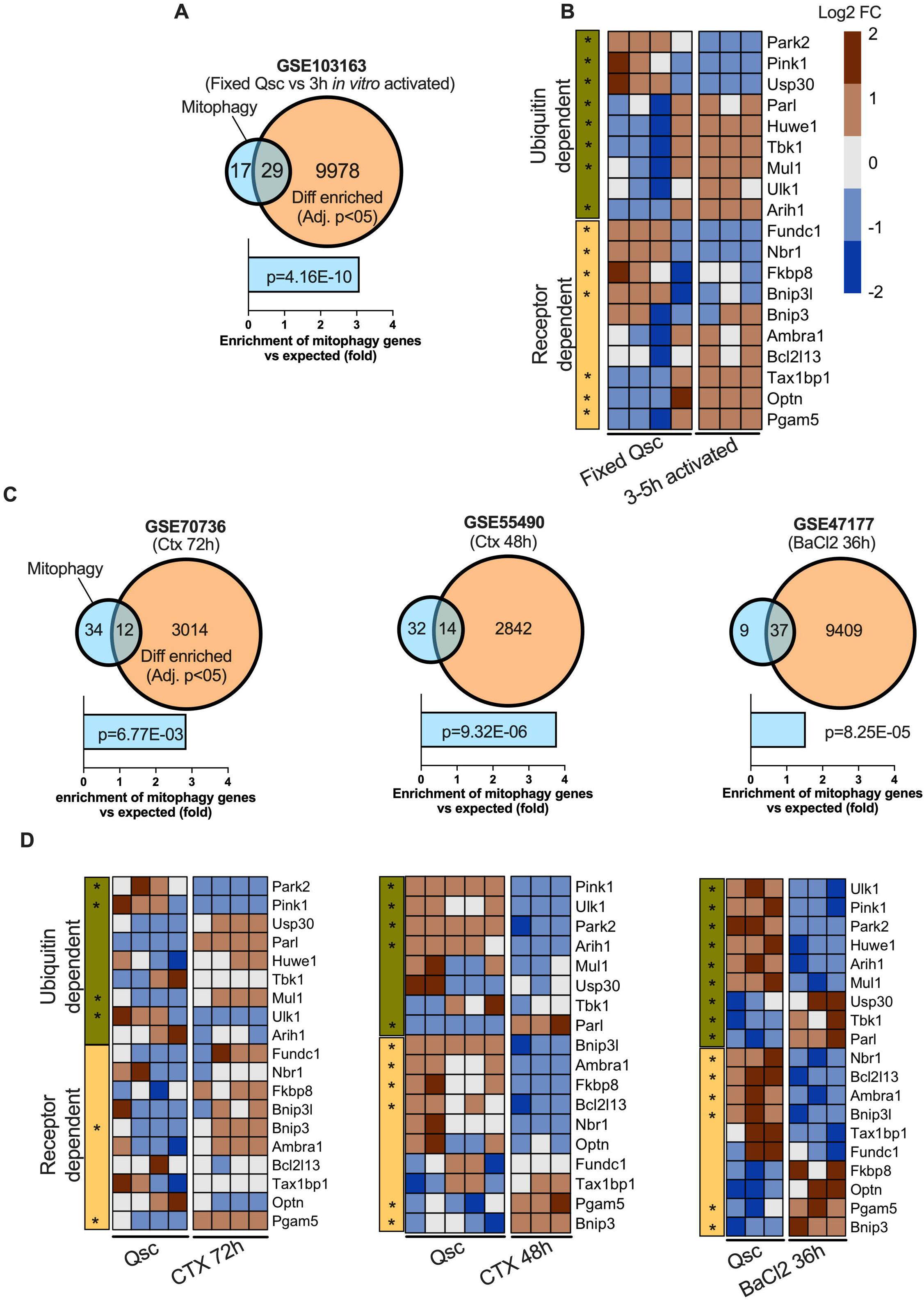
Mitophagy pathways undergo rapid transcriptional changes in MuSCs transitioning from a quiescent to activated state. Four transcriptomics datasets we used to examine transcriptional changes in mitophagy related genes in quiescent and activated MuSCs. One dataset (Panel **A-B**; GSE103162) compared MuSCs from muscles that were fixed prior to enzymatic dissociation to preserves the native quiescent state of cells, to unfixed MuSCs that had undergone activation *in vitro* for 3-5h. The three other datasets (Panel **C-D**; GSE70736, GSE55490, GSE47177 ^14,19,20^) compared quiescent and *in vivo* activated MuSCs 36-72h following muscle injury with Cardiotoxin or Barium Chloride. Venn Diagrams present the number of mitophagy related genes (GO:0000423; 1901524;1903146;0000422) that are differently enriched the datasets. Fold enrichment of mitophagy genes is shown below each Venn diagram along with the hypergeometric test p value. For each dataset, the heatmap shows changes in the expression of genes related to ubiquitin and receptor-dependent mitophagy ranked according to fold change *vs* the quiescent state. Statistical significance (*: q value < 0.05) is shown for each gene.

To examine whether functional differences occurred under these states, mitophagy was assessed in MuSCs residing on EDL myofibers isolated from the mitoQC transgenic mouse line expressing a pH-sensitive mCherry-GFP tag targeted to mitochondria, which allows to identify mitochondria located in acidic lysosomes based on the pH-dependent quenching of GFP fluorescence ^21^ (Fig. 2A-B). MuSCs were examined at 1 and 24h post isolation at which time a majority of cells were in the quiescent and activated state, respectively, based on Pax7 and MyoD staining (Fig. 2 C-D). Results revealed that the number of mCherry^+^ puncta was higher at 1h compared to 24h (Fig. 2E), suggesting that mitophagy is active in quiescent cells, and downregulated in activated cells.

**Figure 2:**
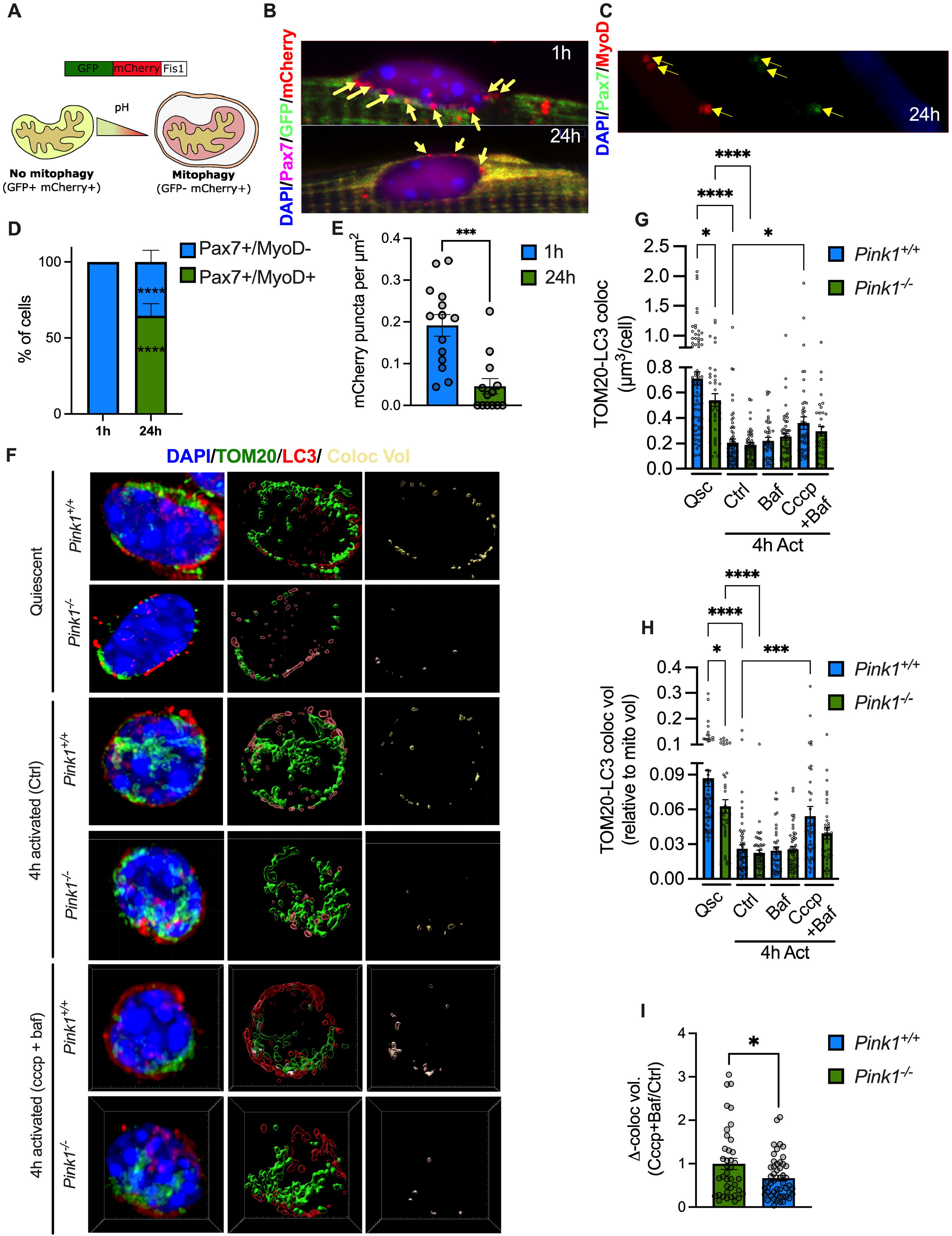
Mitophagy is prominent in quiescent MuSCs and downregulated during activation. **A)** Schematic illustrating how the mCherry-GFP tag targeted to mitochondria allows to identify mitochondria within acidic lysosomes based on the pH-dependent quenching of GFP fluorescence. **B)** Representative images of MuSCs at the surface of EDL myofibers 1 and 24 hours after their isolation from mitoQC mice expressing the mCherry-GFP tag. **C-D)** Proportion of quiescent (Pax7+/MyoD-) and activated MuSCs (Pax7+/MyoD+ and Pax7-/MyoD+) at 1 and 24 hours post isolation (*n*=38-40 MuSC clusters from 8-10 fibers isolated from 2 mice). Representative image is of doubly labeled cells is shown at 24h. **E)** Number of mCherry puncta per μm^2^ of MuSC surface (*n*=13-14 fibers, each containing 4-6 MuSCs) in MuSCs 1 and 24h post isolation. **F)** 3D reconstruction of TOM20-labeled mitochondria (green) and LC3-labeled autophagosomes (red) in freshly sorted quiescent and 4h *in vitro* activated MuSCs isolated from *Pink1*^*+/+*^ and *Pink1*^*-/-*^ mice. The volume of mitochondria overlapping with autophagosomes is shown in yellow. In the right-end panels, mitochondria and autophagosomes have been removed to highlight changes in colocalization. **G-H)** Volume of mitochondria colocalized to autophagosomes expressed in absolute value or relative to total mitochondrial volume (*n*=50-61 cells from 3 mice in each group and experimental condition). **I)** Impact of CCCP on mitochondrial colocalization to autophagosomes expressed as fold change relative to control condition. All data are presented as mean ± sem. *: p<0.05, ***: p<0.001, ****: p<0001 on unpaired two-tailed *t* tests or one way ANOVAs.

### 2.2 Pink1 maintains mitophagy in the quiescent state

Given the results of our transcriptomics analysis showing the consistent enrichment of *Pink1* and *Parkin* transcripts found in the quiescent compared to the activated state, we reasoned that this pathway could be involved in the regulation of mitophagy. To test this, MuSCs were purified from uninjured wild-type and *Pink1*-deficient mouse muscles and mitophagy was assessed in freshly isolated quiescent and *in vitro* activated cells using high resolution confocal microscopy and 3D reconstruction of TOM20-labeled mitochondria and LC3^+^ autophagosomes (Fig. 2F). Activation was allowed for only 4 hours to examine whether changes in mitophagy occurred rapidly during state transition. In freshly isolated quiescent cells, colocalization of mitochondria and LC3^+^ autophagosomes was prominent, representing 6-9% of cellular mitochondrial content (Fig. 2F-H). Interestingly, during the 4 hour activation period, a substantial reduction of colocalization occurred (Fig. 2F-H). A similar pattern was also observed when colocalization between mitochondria and lysosomes was quantified (Fig. S1), which was altogether consistent with changes in mitophagy observed in mitoQC EDLs during transition from quiescence to activation. In PINK1 deficient cells, mitochondrial colocalization to autophagosomes (Fig 2F-H) and lysosomes (Fig. S1) was mildly reduced, under the quiescent state, but not in 4h-activated cells, presumably because of the low levels of mitophagy observed in the latter condition.

To determine whether loss of colocalization during transition to the activated state was due to reduced mitophagy or increased lysosomal activity, Bafilomycin-A was added during the 4h activation period to block lysosomal acidification and fusion with autophagosomes (Fig. 2F-H). Irrespective of the genotype, Bafilomycin-A had a negligible impact on colocalization, confirming that mitophagy is low in 4h activated MuSCs. Interestingly, addition of CCCP in presence of Bafilomycin-A, was able to increase colocalization between mitochondria and autophagosomes, suggesting that MuSCs can react to a potent mitophagy trigger (Fig. 2I). However, in absence of PINK1, the increase in colocalization observed in presence of CCCP was reduced compared to *Pink*^*+/+*^ controls (Fig. 2I), suggesting a reduced ability to activate depolarization-dependent mitophagy. Altogether these results suggested that PINK1 contributes to maintain mitophagy in the quiescent state.

### 2.3 Loss of Pink1 promotes MuSC commitment at the expense of self-renewal leading to a progressive loss of muscle regenerative capacity

Considering the differences in mitophagy observed between the quiescent and the activated state, we reasoned that PINK1 deficiency could affect fate decisions. To test this, MuSC state was tracked in EDL fibers cultured for up to 72h. Activation was more rapid in *Pink1*^*-/-*^ MuSCs compared to controls as indicated by a higher proportion of Pax7^+^/MyoD^+^ cells at 24h in culture (Fig. 3A-B). Importantly, by 72h *Pink1*^*-/-*^ MuSCs exhibited a higher propensity to commit to the myogenic lineage at the expense of self-renewal as evidenced by an increased proportion of Pax7^-^/MyoD^+^ cells, and a reduced proportion of cells expressing Pax7 only (Fig. 3A-B). No differences in the number of MuSCs per fiber at baseline (1h) (Fig. 3C) or the number of cells per cluster of dividing MuSCs (72h) (Fig. 3D) were observed between *Pink1*^*-/-*^ and controls indicating no gross effect of PINK1 deficiency on MuSC viability and proliferation.

**Figure 3:**
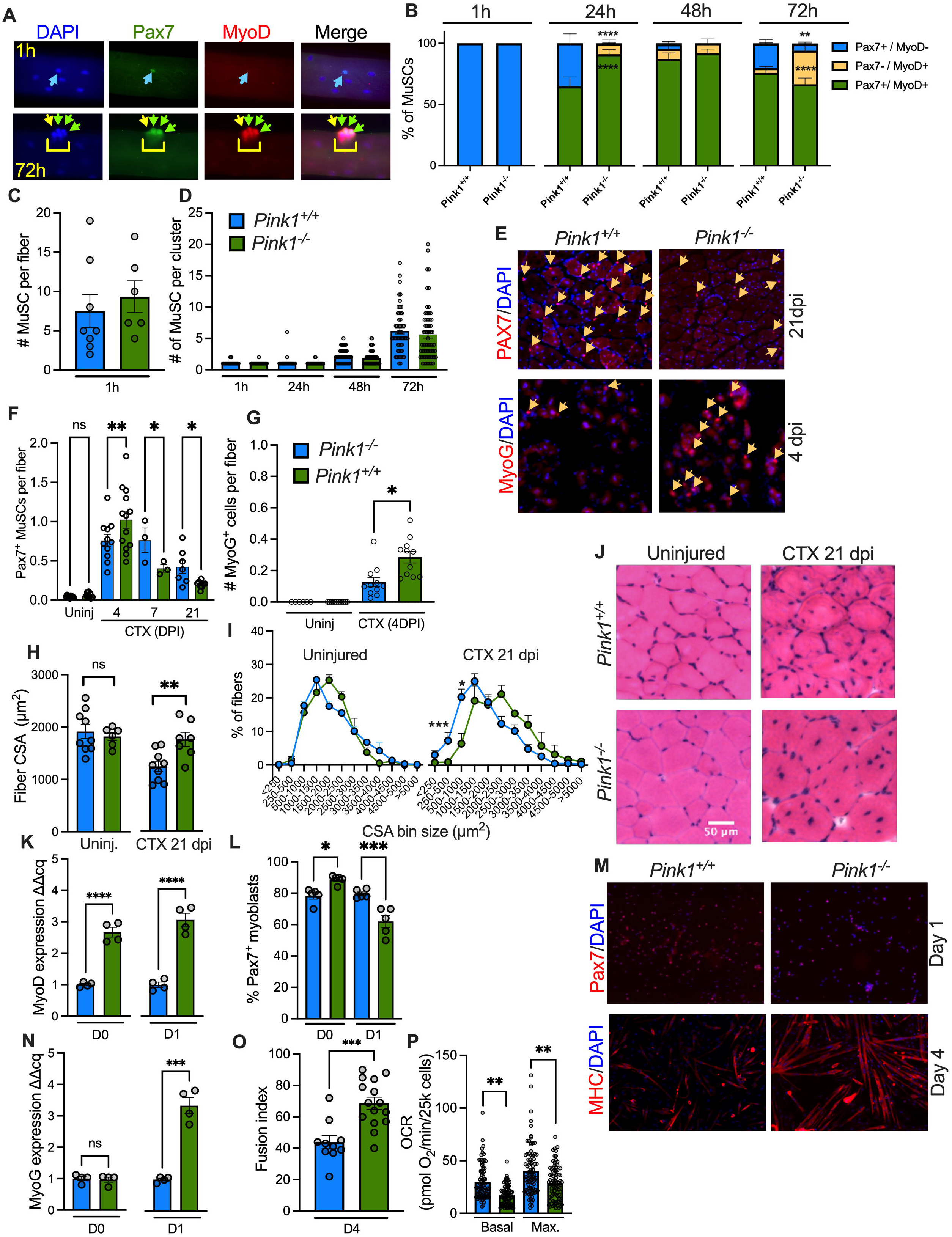
Pink1 deficiency impairs MuSC fate decision but has no adverse impact on differentiation. **A)** Representative images of Pax7 and MyoD labeled MuSCs at the surface of EDL fibers at after 1 and 72h of culture. **B)** Proportion of quiescent (Pax7^+^/MyoD^-^) and activated MuSCs (Pax7^*+*^/MyoD^+^ and Pax7^-^/MyoD^+^) on EDL myofibers at the indicated time points (*n*=45-65 MuSC cluster on 3-5 fibers isolated from 3 mice in each group).**C)** Number of Pax7^+^ cells per EDL myofiber 1h after isolation (each dot represent the average of 3-5 fibers per well isolated from 3 mice). **D)** Number of cells per MuSCs cluster at the indicated time points (*n*=45-65 MuSC cluster on 3-5 fibers isolated from 3 mice in each group). **E)** Representative images of Pax7^+^ and MyoG^+^/DAPI stained TA cross sections at 21 and 4 days post CTX injury. **F-G)** Number of Pax7^+^ and MyoG positive cells in uninjured and injured TA muscles at various time points following CTX injection. Given the differences observed in fiber CSA between genotypes, cell count is expressed as the number of Pax7^+^ or MyoG^+^ cells divided by the number of fibers present in each 500 μm^2^ ROI analyzed (*n*=9-12 ROIs from 3-4 TA muscle per group). **H-J)** Representative H&E stains, mean cross sectional area (CSA) and size distribution of fibers in contralateral uninjured and injured TA muscle of *Pink1*^*+/+*^ and *Pink1*^*-/-*^ mice 21 days after CTC injury. **K)** MyoD transcript levels at d0 and d1 into differentiation (*n*=4 distinct RNA isolations). **L)** Proportion of Pax7^+^ myoblasts at d0 and d1 into differentiation (*n*=5 distinct culture wells). **M)** Representative images of Pax7/DAPI and eMHC/DAPI stained myoblasts from *Pink1*^*+/+*^ and *Pink1*^*-/-*^ mice at d1 and d4 into and differentiation. **N)** MyoG transcript levels at d0 and d1 into differentiation (*n*=4 distinct RNA isolations). **O)** Proportion of nuclei in eMHC^+^ polynucleated myotubes (*i*.*e*. fusion index) measured at d4 into differentiation (*n*=10 distinct culture wells). **P)** Basal and maximal CCCP (1μM) stimulated oxygen consumption rate (OCR) in *Pink1*^*+/+*^ and *Pink1*^*-/-*^ myoblasts (*n*=78-84 wells, 3 distinct cell cultures per genotype). *: p<0.05, **: p<0.01, ***: p<001 ****: p<0.0001 on unpaired two-tailed *t* tests or one way ANOVAs.

To assess whether fate decision was dysregulated *in vivo*, TA muscle were injured with CTX and MuSC state was examined at 4 and 21 days post-injury to capture the proliferation/commitment, and the late regenerative phases. As shown in Fig. 3E-F, the number of Pax7^+^ MuSCs was higher in *Pink*^*-/-*^ compared to control muscles at 4 days, but this was reversed at 21 days post injury at which time the number of Pax7 cells was ∼50% lower compared to *Pink1*^*+/+*^ controls. Furthermore, the number of cells expressing MyoG^+^ was increased in *Pink*^*-/-*^ mice at 4 days (Fig. 3E, G), which altogether confirmed that in absence of PINK1, MuSCs display premature commitment at the expense of self-renewal, leading to a reduced MuSC pool size in regenerated muscle.

Histological assessment of TA muscles at 21 days post injury revealed that regeneration was more robust in *Pink1*^-/-^ mice and yielded fibers with a larger cross sectional area (CSA) (Fig. 3H-J). This suggested that the capacity to differentiate was not adversely affected in absence of PINK1. To test this directly, primary myoblasts were cultured under proliferating conditions and subsequently switched to differentiation media. Transcript levels for MyoD were higher in PINK1 deficient myoblasts (Fig. 3.K). Furthermore, after switching to differentiation media, the proportion of Pax7^+^ cells declined more readily (Fig. 3L-M), and the expression of MyoG turned-on more rapidly in *Pink1*^*-/-*^ myoblasts (Fig. 3N). Moreover, the formation of MHC^+^ myotubes was increased in absence of PINK1 (Fig. 3M, O). Together, these results indicated that differentiation capacity is not only preserved but enhanced in absence of PINK1 due to increased commitment. Interestingly, baseline and maximal rates of oxygen consumption were reduced in PINK1 deficient myoblasts (Fig. 3P), but these bioenergetic abnormalities clearly had no adverse impact on the capacity to differentiate.

To assess the importance of PINK1-dependent mitophagy in maintaining MuSC regenerative capacity in the longer term, mice were exposed to repeated CTX injuries each separated by 21 days. Regenerated fibers became smaller in *Pink1*^*-/-*^ mice (Fig. 4A-C), and the number of *Pax7*^*+*^MuSCs declined further (Fig. 4D-E) after repeated cycles of injury, indicating that altered fate decision in absence of PINK1 leads to a progressive loss of muscle regenerative capacity.

**Figure 4:**
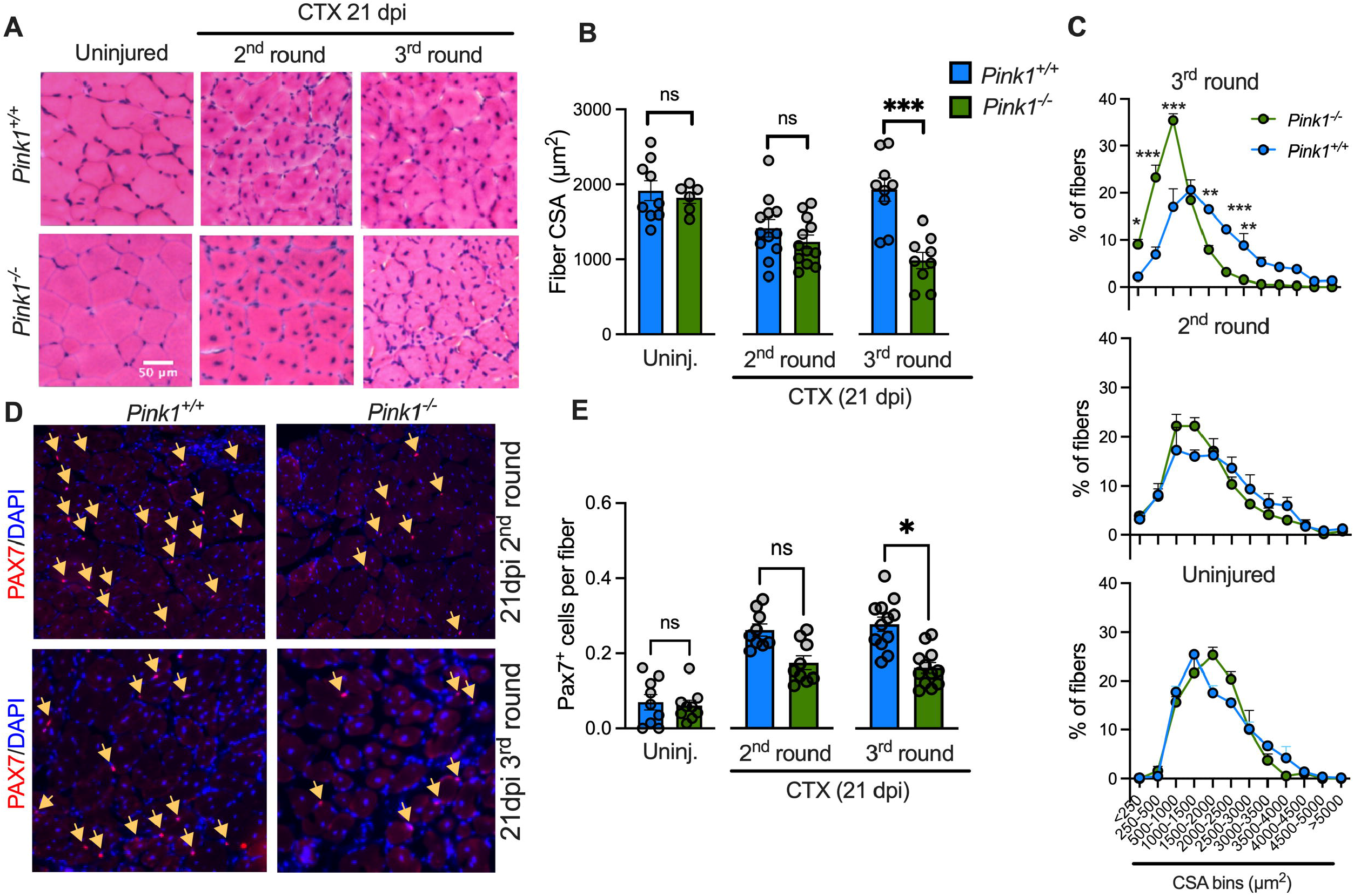
Pink1 deficiency leads to a progressive loss of muscle regenerative capacity.. Representative H&E stains (**A**), mean cross sectional area (CSA) (**B**) and size distribution (**C**) of fibers in contralateral uninjured and injured TA muscle of *Pink1*^*+/+*^ and *Pink1*^*-/-*^ mice 21 days after a 2^nd^ and 3^rd^ cycle of CTX injury-regeneration (*n*=9-12 ROIs from 3-4 TA muscle per group). **D)** Representative images of Pax7/DAPI stained TA cross sections 21 days after a 2^nd^ and 3^rd^ cycle of CTX injury-regeneration. E) Number of Pax7^+^ positive cells in uninjured and injured TA muscles after the a 2^nd^ and 3^rd^ cycle of CTX injury-regeneration. Given the differences observed in fiber CSA, MuSC pool size is expressed as the number of Pax7^+^ cells divided by the number of fibers present in each 500 μm^2^ ROI analyzed (*n*=9-12 ROIs from 3-4 TA muscle per group). *: p<0.05, **: p<0.01, ***: p<001 on unpaired two-tailed *t* tests or one way ANOVAs.

### 2.4 Impaired fate decisions in Pink1 deficient MuSCs is linked to increased mitochondrial ROS generation

Previous studies showed that ROS release is one of the key signaling factors through which mitochondria can orient stem cells toward self-renewal or commitment ^12,22,23^. We therefore suspected that increased mitochondrial ROS release could underlie impaired fate decisions in the absence of PINK1. To test this, mitochondrial superoxide release was measured in MuSCs at the surface of freshly isolated EDL fibers. Mitochondrial superoxide release, measured using MitoSOX, increased by 36% in PINK1 deficient MuSCs compared to wild type controls (Fig. 5A-B). However, no significant differences between the two genotypes were observed when the complex I inhibitor rotenone was added to maximize mitochondrial superoxide production (Fig. 5A-B). Experiments performed in permeabilized cells revealed that mitochondrial ROS release resulting from PINK1 deficiency persisted at the myoblast stage as evidenced by an increased release of H_2_O_2_ in presence of substrates feeding the electron transport chain at complex I and II (Fig. 5C).

**Figure 5:**
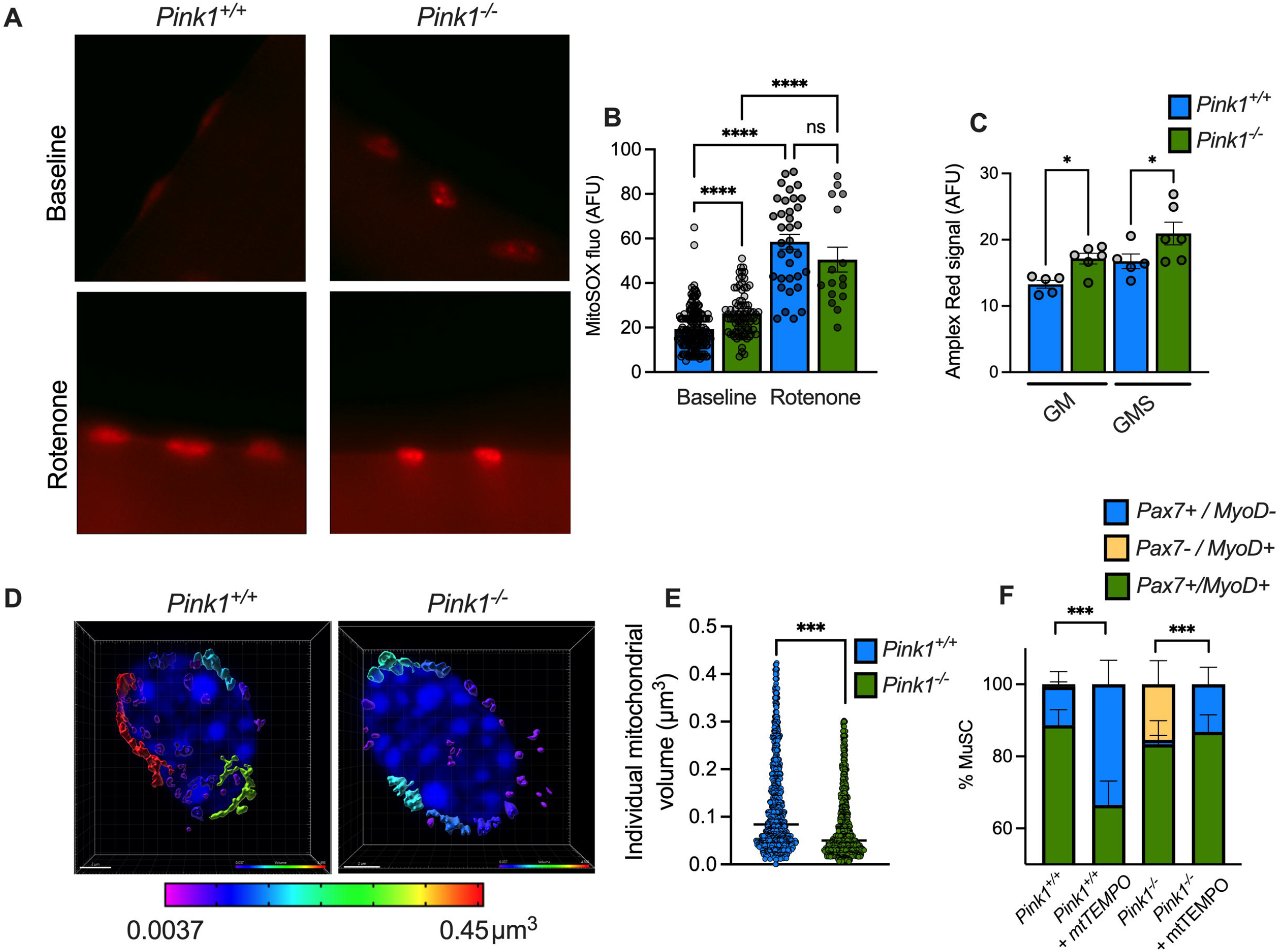
Impaired fate decisions in Pink1 deficient MuSCs is linked increased mitochondrial ROS generation. Representative images (**A**) and (**B**) quantification of mitoSOX fluorescence in MuSCs at the surface of EDL fibers isolated from *Pink1*^*+/+*^ and *Pink1*^*-/-*^ mice (*n*=80-170 cells, 16-30 fibers, 3 mice per genotype). Measurements were performed in absence or presence of 1 μM rotenone. **C)** Mitochondrial H_2_O_2_ release measured by Amplex Red fluorescence in permeabilized myoblasts following successive addition of glutamate-malate (5:2.5 mM), and succinate (5 mM) (*n*=5-6 replicates from 3 distinct cell cultures per genotype). **D-E)** Volume of individual mitochondria in quiescent MuSCs from *Pink1*^*+/+*^ and *Pink1*^*-/-*^ mice (*n*=1130-1227 mitochondria in 33 cells isolated from 3 mice in each group). **F)** Proportion of quiescent (Pax7^+^/MyoD^-^) and activated MuSCs (Pax7^+^/MyoD^+^ and Pax7^-^/MyoD^+^) in EDL myofibers from *Pink1*^*+/+*^ and *Pink1*^-/-^ mice cultured in absence or presence of 10 μM MitoTEMPO for 24h (*n*=78-90 MuSCs, from 5 fibers isolated from 4 mice in each genotype). *: p<0.05, ***: p<001 ****: p<0.0001 on a two-tailed unpaired t test or one way ANOVA.

Since mitochondrial dynamics was shown to regulate stem cell fate through ROS signaling, the state of the mitochondrial network was also examined in quiescent MuSCs. Results indicated a significant reduction in the average volume of individual mitochondria in *Pink1*^*-/-*^ MuSC, indicating a more fragmented mitochondrial network (Fig. 5D-E). Altogether, these results indicated that in absence of PINK1 mitochondria are in a configuration that promotes increased ROS production.

EDL myofibers from control and *Pink1*^*-/-*^ mice were thus cultured for 24 hours in presence or absence of the mitochondria-targeted superoxide scavenger mitoTEMPO to assess the impact on fate decisions. As shown in Fig. 5F, mitoTEMPO reduced the proportion of committed (Pax7^-^/MyoD^+^) cells in *Pink1*^*-/-*^ mice, and restored the proportion self-renewing (Pax7^+^/MyoD^-^) cells to the levels observed in untreated wild type mice, confirming that increased ROS production is an important factor underlying impaired fate decision in absence of PINK1.

## 3 DISCUSSION

Maintenance of optimal mitochondrial function plays a crucial role in the regulation of muscle MuSC behavior, including their decision to self-renew or commit and differentiate ^4–7^. However, the role of specific autophagy mechanisms such as mitophagy in regulating mitochondrial qualities and MuSC states remains largely unknown. Results from the present study show that mitophagy is prominent during quiescence and is rapidly decreased upon activation, pointing to a role in regulating MuSC state transitions. Importantly, our data also reveal that while PINK1 is dispensable for normal differentiation to proceed, it is required for normal mitophagy to occur in the quiescent state, which allows to maintain MuSC self-renewal by limiting mitochondrial ROS production. Mild impairment of mitophagy in absence of PINK1 is therefore sufficient to disrupt MuSC fate decision ultimately leading to a loss of regenerative capacity.

Quiescent stem cells, including MuSCs contain a relatively small number of mitochondria arranged in low complexity networks ^11,12,22,23^. Their rate of oxidative phosphorylation is low and dependent on fatty acid metabolism ^9,11,12,24^, while mitochondrial membrane potential^24^ and ROS release ^12,22,23,25^ are reduced. The current view is that this configuration allows mitochondria to emit pro-quiescence signals, while limiting oxidative damage^4,6,7,26^. The prominence of mitophagy in quiescence, and the loss of PINK1 blunting mitophagy, enhancing mitochondrial ROS release, and impairing self-renewal unveil an important role for PINK1/Parkin dependent mitophagy in maintaining the metabolic configuration of quiescence/self-renewal. Although our data indicate that protection against excess mitochondrial ROS emission is a key factor by which mitophagy promotes MuSC quiescence/self-renewal, mitochondrial culling may also help preserve other mitochondrial traits that contribute to maintain stemness through nutrient sensing signaling pathways, epigenetic, or post-translational protein modifications ^4,6,7,26^.

Because of their low metabolic rate, stem cells are not particularly prone to wear and tear. It therefore remains unclear why a pathway such as mitophagy, which is commonly depicted as a mechanism to clear damaged/dysfunctional organelles, would be prominent in the quiescent state. Considering that mitochondrial depolarization is a well-established trigger for PINK1-mediated mitophagy ^27–29^, one possibility could be that the low membrane potential observed in quiescent stem cells ^10,24^ poises mitochondria toward mitophagy compared to more metabolically active cells. If true, PINK1-dependent mitophagy could thus be viewed as a preventive mechanism which, if defective or physiologically inhibited, promotes the emergence of polarized, ROS emitting mitochondria that reduce quiescence depth, and pushes MuSCs toward activation and commitment.

While mitophagy is required to maintain MuSC self-renewal, previous studies in cultured myoblasts have reported that upregulation of autophagy/mitophagy also plays a crucial role in supporting the metabolic/mitochondrial remodeling process required for proper myoblast differentiation, although the mitophagy pathways involved remain poorly defined ^30,31^. Our result showing that *in vitro* and *in vivo* differentiation capacity is preserved in PINK1 deficient myoblasts clearly indicate that mechanisms other than the PINK1/Parkin pathway are involved. In this regard, our transcriptomics analysis showed that the expression of *Pink1* and *Parkin* was consistently high in quiescent MuSCs, and that the repression of their expression during activation was accompanied by an upregulation of several genes involved in receptor-dependent mitophagy, including the mitophagy receptor *Bnip3*. Interestingly, previous studies in c2c12 myoblasts have shown that inactivation of *Bnip3* disrupts differentiation and impairs mitochondrial remodeling, as evidenced by blunted changes in the abundance of several mitochondrial marker proteins ^30^. Collectively, these result shed new light on the regulation of mitophagy during myogenesis and suggest that various pathways come into play at various stages, with PINK1-dependent mitophagy being mostly important for quality control in MuSCs, as shown in the present study, while BNIP3 and potentially other mitophagy receptors could predominate during differentiation.

## 4 METHODS

### 4.1 Animal care

*Pink1* (PTEN induced putative kinase 1) knockout mice were obtained from Dr J. Shen (strain B6.129S4-*Pink1*^*tm1Shn/J*^) and were backcrossed to C57BL/6J for more than seven generation, as described in ^32^. Mitophagy reporter mice (*Mito*-QC) mice harboring a mitochondria-targeted tandem GFP-mCherry reporter were obtained from Dr. Ian Ganley^21^. All experiments on animals were approved by the University of Ottawa Institutional Animal Care Committee and conducted according to the directives of the Canadian Council on Animal Care. Mice were maintained in ventilated cage racks by groups of 3-5 mice. All mice of both sexes were kept on a regular 12–12 h light-dark cycle, and had access to food and water *ad libitum*. Animals were used at 8-12 weeks of age for experiments and euthanized by cervical dislocation.

### 4.2 Cardiotoxin injury

Mice were injected subcutaneously with Buprenorphine (0.1 mg/Kg) and anesthetized by gas inhalation 30 min prior to Cardiotoxin (CTX) injection. CTX diluted in PBS (50 μL of 10 μM) was injected into the *Tibialis Anterior* (TA) muscle and the uninjected contralateral muscle was used as control. Mice recovered in a cage with a heating pad and were monitored 24 hrs post-CTX injection. This procedure was repeated up to three time every 21 days to test MuSC regenerative potential following successive rounds of injury-regeneration.

### 4.3 Muscle tissue preparation

At selected time points following CTX injection, TA muscles were dissected, weighed and cut in half in a cross-sectional orientation. One portion was embedded in Optimal Cutting Temperature (OCT) compound and flash frozen in cooled isopentane, while the other portion was dropped in freshly prepared paraformaldehyde (2% w/v, 4 °C) and fixed for 30 minutes. Following fixation, TA muscles were washed twice with PBS for 5 minutes, twice with glycine (0.25M in PBS) for 10 minutes, and then left in 5% (w/v) sucrose for 2 hours, followed by 20% sucrose (w/v) for 2-3 days. TA muscles were then embedded in OCT compound, frozen in cooled isopentane and stored at -80°C. Tissues were cryo-sectioned in a cross-sectional orientation at a thickness of 14 μm and glass slides were stored at -80°C until further processing.

### 4.4 Single EDL fiber isolation and culture

The extensor digitorum longus (EDL) muscles were harvested immediately following cervical dislocation and digested in 0.5% (w/v) collagenase B for approximately 30-40 minutes at 37°C. Muscles were transferred in wash media (DMEM containing 4.5 g/L glucose and 1% (v/v) penicillin-streptomycin), and single fibers were obtained by gentle trituration of muscles with a wide bore pipette. Single fibers were then freed from debris through two successive washes. Following a 10 min rest period, fibers where either fixed in warmed PFA (2% w/v) for 10 min to capture the quiescent state as recently described^12^. Some fibers were cultured over 12-72h (in DMEM 4.5 g/L glucose, 20% (v/v) FBS, 1% (v/v) chicken embryo extract, 1% (v/v) penicillin-streptomycin, 7.5 ng/mL βFGF) prior to fixation in order to track MuSC activation and commitment. Fixed fibers were kept at 4 °C until further processing.

### 4.5 Muscle stem cell purification and culture

Hindlimb muscles were harvested, pooled, and digested in 1% (w/v) Collagenase-B and 0.4% (w/v) Dispase II in Hams-F10 media, using the Milteny MACS Octo-dissociator SLICE_FACS program for 27 minutes as per^12^. Following digestion, the muscle slurry was filtered (100μm mesh size) and centrifuged (10 min, 500xg). The cell pellet was resuspended in red blood cell lysis buffer and incubated for 5 minutes to remove residual erythrocytes. Cells we then spun down (5 min, 500xg) and washed in FACS buffer (5mM EDTA, 10% (v/v) FBS in 1xPBS) before being resuspended in 1 mL of FACS buffer containing the following antibodies: A) PE-conjugated Sca-1, CD45, CD31 and CD11b to remove non-MuSCs present in the population, and B) 647-conjugated α integrin-7 and VCAM-1 to positively select α-integrin-7 ^+^/VCAM^+^ MuSCs. Following incubation with antibodies (15-25 min at 4°C), the volume was topped to 15 mL with FACS buffer and cells were centrifuged 5 min at 500xg. The resulting pellet was resuspended in 1 mL of FACS buffer, filtered through a 50 μm CellTrics filter into a 5 mL round bottom tube. Cells were then sorted by FACS at the Flow Cytometry facility at the Ottawa Hospital Research Institute (OHRI).

### 4.6 Primary myoblast isolation and culture

Myoblasts were prepared from skeletal muscle as per ^33^. Hindlimb muscles were harvested then minced in a 1% (w/v) Collagenase-B and 0.4% (w/v) Dispase II solution in Hams-F10 media. The tissue was then digested for 30 minutes at 37°C and mixed at the halfway point. The slurry was diluted in 25ml Hams-f10 media and filtered through a 70μm filter. The filtered solution was centrifuged (100g 5 minutes) and the pellet resuspended in Myoblast Culture Media (Hams-F10 Media, 20% FBS, 1% Penicillin-Streptomycin and 2.5ng/ml Basic Fibroblast Growth Factor, human (hbFGF)). The cell suspension was pre-plated on a non-coated tissue culture plate and after 1.5 hours transferred to a collagen coated plate (0.01% collagen Type 1, 0.2% Acetic acid in ddH20 and Filter sterilized). Myoblasts were further filtered by pre-plating until all contaminating cells were removed.

Myoblasts were cultured in Myoblast Culture Media (changed every second day) until around 70% confluent when they were split. Splitting was performed by washing 3x in PBS in a 20% Tryspin/EDTA (0.25% trypsin/ 2.21mM EDTA 4Na) PBS solution at 37°C until detached. Myoblast differentiation was performed by switching to a Differentiation Media (DMEM containing 4.5 g/L glucose 2% FBS and 1% Penicillin-Streptomycin) changed daily for up to fours days.

### 4.7 Hematoxylin & Eosin staining

Cross sections from flash frozen samples were stained with Hematoxylin and Eosin (H&E) at the uOttawa Histology Core Facility. Images were taken at 20x magnification on an EVOS Fl Auto2 microscope and digitally stitched to reconstitute entire muscle sections. For each muscle, myofiber numbers and minimum Ferret diameter were measured on three 1000 μm^2^ regions of interest (ROI) using Fiji (ImageJ). Myofiber cross sectional area (CSA) was calculated using the equation CSA = π x (minimum Ferret diameter/2)^2^.

### 4.8 Immunofluorescence

#### 4.8.1 Muscle sections

Tissue sections were submitted to antigen retrieval in citrate buffer (10 mM sodium citrate, 0.05% Tween-20, pH=6.0), followed by 1 hour blocking in 3% (w/v) BSA. Slides were incubated overnight at 4°C with primary antibodies against Pax7, MyoD, MyoG, alone or together with Ki67. Slides were then washed 5 times with 1xPBS, and fluorescence-conjugated secondary antibodies, (AlexaFluoro 488, AlexaFluoro 594 or CY3) diluted in 3% BSA were added to sections along with DAPI. Following incubation at room temperature for 15 min, sections were washed 5 times and slides were mounted using Prolong Gold with DAPI. Images were taken at 20x (0.4 NA) magnification on an EVOS Fl Auto2 microscope, digitally stitched to reconstitute entire muscle sections, and analyzed using Fiji (ImageJ). For each muscle section, the number of MuSCs was counted in three distinct 1000 μm^2^ ROIs and normalized by the number of muscle fibers present in each ROI.

#### 4.8.2 Fate determination in EDL fibers

Single EDL myofibers were permeabilized in 0.1% (v/v) Triton-X and 100 mM glycine, followed by blocking in blocking solution (5% (v/v) horse serum, 2% (w/v) BSA and 0.1% (v/v) Triton-X in PBS) for 5 hours at room temperature. Myofibers were incubated overnight at 4 °C with a combination of the following primary antibodies diluted in blocking solution: Pax7, MyoD, and MyoG. Following incubation with secondary antibodies (AlexaFluoro 488, AlexaFluoro 594) for 1 hour at room temperature, myofibers were mounted on charged glass slides using Immu-mount and imaged at 100x oil immersion (1.28 NA) magnification on an EVOS Fl Auto2 microscope. Each myofiber was manually inspected throughout its thickness and along its full length to capture the entire population of MuSC.

#### 4.8.3 Purified MuSCs and cultured myoblasts

Cells were fixed in 4% PFA, washed 3X in PBS and quenched in 50mM NH_4_Cl. Cells were permeabilized in 0.1% (v/v) Triton X-100 and washed with PBS. Cells were blocked in 10% (v/v) FBS for 1 hour and incubated with primary antibodies in 5% FBS overnight. Primary antibodies included Pax7, MHC, TOM20, LC3 and LAMP1. Cells were incubated with secondary antibodies (AlexaFluoro 488, AlexaFluoro 594), then incubated with DAPI 300nM for 3 minutes. Cells were then washed, cover slipped, and imaged either on an EVOS Fl Auto2 microscope (20X (0.4 NA) magnification) to quantify myoblast differentiation, or a Zeiss LSM880 AxioObserver Z1 (63X magnification 1.4 NA, Oil, Plan-Apo) to quantify mitophagy and mitochondrial morphology.

### 4.9 Myoblast differentiation

To assess differentiation, the fusion index, representing the proportion of DAPI-stained nuclei incorporated in multi-nucleated MHC-positive myotubes was quantified four days after switching to differentiation media using the Fiji (ImageJ).

### 4.10 Confocal image processing and quantification of mitophagy

To analyze mitochondrial morphology and colocalization to either autophagosomes or lysosomes, cells labeled with TOM20 and either LC3 or LAMP1 were imaged using high resolution confocal microscopy and processed using the Zeiss Aryscan processing function. To control for batch effects and sampling biases, cells from *Pink1*^*+/+*^ and *Pink*^*-/-*^ mice were always isolated and processed in parallel for each experimental day, and multiple cells were randomly imaged in each experiment. To control for image processing and quantification biases, 3D-reconstruction and calculation of colocalization volume was batch processed in the IMARIS software using fixed settings for the intensity and voxel size threshold values, and surface/surface colocalization parameters.

### 4.11 RNA isolation and qPCR

RNA was isolated from primary myoblasts using the Quigen RNeasy kit according to the manufactures instructions. 1000ng of RNA from isolations were transformed to cDNA by using the BioRad iScript™ cDNA Synthesis Kit. qPCR was performed using the BioRad iTaq Universal SYBR Green Supermix. Each sample was run in triplicate using the following thermal cycler programming: 95°C for 2 minutes,, cycling of 95 °C for 5 seconds and 60 °C for 30 seconds for 40 repeat cycles, a melt curve was run from 65-95 °C with a 0.5°C rise each step qPCR runs were performed alongside a housekeeping gene (TBP and/or Hkpt2) for normalization.

### 4.12 Respirometry

Oxygen consumption rates (OCRs) were measured in myoblasts using a Seahorse extracellular flux analyzer. 3×10^4^ cells were plated in XF96 well plates in basal XF media. Baseline OCR was recorded for 20 min followed by the sequential addition of oligomycin (1μM), CCCP (2 μM) and Antimycin-A/Rotenone (0.5 μM). Cells were then labeled with Hoescht 33342 (1 μM), and wells were imaged on an EVOS Fl Auto2 microscope. Cell counting was performed using the Cell Profiler software and values were reported in pmol O_2_ min ^−1^ 25,000 cells.

### 4.13 Determination of mitochondrial ROS release

For the analysis of mitochondrial superoxide release in MuSCs, single EDL myofibers were isolated as described, and incubated in culture media supplemented with mitoSOX Red (5μM, 15 min, 37 °C) in absence and presence of 1 μM rotenone. Myofibers where then washed in 1xPBS and mounted in charged glass slides. MuSCs at the surface of myofibers were imaged at 100x using an oil immersion objective, and fluorescence was analyzed on Fiji (ImageJ). A ROI was drawn around each MuSC and background fluorescence intensity subtraction was performed using a 50 pixels rolling ball radius. Data was analyzed as maximal fluorescence intensity within each ROI.

For the measurement of mitochondrial H_2_O_2_ release 5×10^6^ myoblasts were permeabilized with digitonin (2μg/μl) for 2.5 min and immediately incubated at 30°C under continuous stirring in 600 μl of buffer Z (110mM K-MES, 35mM KCl, 1mM EGTA, 5mM K2HPO4, 3mM MgCl26H2O, and 0.5 mg/ml BSA, pH 7.3 at 4°C with 100 μM PMSF) supplemented with the H_2_O_2_-sensitive probe Amplex red (1.5 μM;) and horseradish peroxidase (1.2 U/ml). Net H_2_O_2_ release was monitored fluorometrically (ex-em: 563– 587 nm) on a Hitachi F2700 fluorimeter under baseline non-phosphorylating conditions following sequential addition of glutamate-malate (5:2.5 mM) and succinate (5 mM). Rate of H_2_O_2_ release is reported in arbitrary fluorescence units per million cells.

### 4.14 Data mining

The Eulerr R Package was used to generate Venn diagrams illustrating the enrichment of mitophagy related genes in publicly available transcriptomics datasets (GSE103162, GSE70736, GSE55490, GSE47177) ^14,18–20^. For each dataset, the list of differentially expressed genes (q value<0.05) was extracted and compared to a list of mitophagy related genes generated from Gene Ontology terms GO:0000423, GO:1901524, GO:1903146 and GO:0000422. Log2 fold change in the expression of mitophagy-related genes was extracted from the datasets and represented as heatmaps using the pheatmap package for R.

### 4.15 Statistics and reproducibility

For experiments performed in single FACS purified MuSCs, cultured EDL myofibers or myoblasts, a minimum of three different isolations/culture with a minimum of 10 MuSCs or fibers, or 3-10 distinct culture wells was analyzed. For experiments involving *in vivo* CTX injury, a minimum of 3-4 mice per genotype were used and for each TA muscle analyzed 3 distinct ROIs of 1000 μm^2^ each was included to cover a large proportion of the entire muscle. Values are reported as mean ± sem and represented as bar graphs along with individual datapoints when appropriate. Unpaired two-sided t-tests with Welch’s correction (GraphPad Prism 8.4.3) were used to determine statistical difference when two means were compared. To compare more than two means, Brown-Forsythe and Welch ANOVA tests were used with Dunnett’s T3 multiple comparisons test performed for each pair of means compared. For data mining, hypergeometric tests were used to determine the statistical significance of the enrichments of mitophagy related genes in publicly available transcriptomics datasets comparing quiescent and activated MuSCs.

### 4.16 Data availability

Source data used to generate figures are provided as supplemental data. All other data are available from the corresponding authors upon reasonable request.

## 5 FIGURE LEGENDS

**Figure S1: Mitophagy is prominent in quiescent MuSCs and downregulated during activation. A)** 3D reconstruction of TOM20-labeled mitochondria (green) and LAMP1 labeled lysosomes (red) in freshly sorted quiescent and 4h *in vitro* activated MuSCs. The volume of mitochondria overlapping with lysosomes is shown in yellow. In the right-end panels, mitochondria and lysosomes were removed to highlight changes in colocalization. **B)** Volume of mitochondria colocalized to lysosomes under these two state in MuSCs from *Pink1*^*+/+*^ and *Pink1*^-/-^ mice (n=33 MuSCs from 3 mice in each group and experimental condition). *: p<0.05, ***: p<001 on a one way ANOVA.

## 6 AUTHOR CONTRIBUTIONS

Conception of hypothesis: YB. Design of the work: YB, MK, MTM, GC. Acquisition of the data: GC, MTM, MA, NL, AP. Drafting the manuscript: YB, MK, MTM, GC. All authors: Analysis and interpretation of the data and approval of the final version of the manuscript.

## 8 GEOLOCATION INFORMATION

Ottawa, Ontario, Canada

## 9 ACKNOWLEDGEMENTS

The authors gratefully acknowledge services provided by the Louise Pelletier histology core (RRID: SCR_021737) and the Cell Biology and Image Acquisition Core (RRID: SCR_021845) funded by the University of Ottawa, the Natural Sciences and engineering Research Council of Canada, and the Canada Foundation for Innovation. The authors also acknowledge the expert contribution of *Fernando Ortiz from Flow Cytometry & Cell Sorting Facility at OHRI for its contribution on FACS, RRID:SCR_023349*..

