## Supplementary figures and images for "PINK1 Deficiency Alters Muscle Stem Cell Fate Decision and Muscle Regenerative Capacity"

### Figure S1

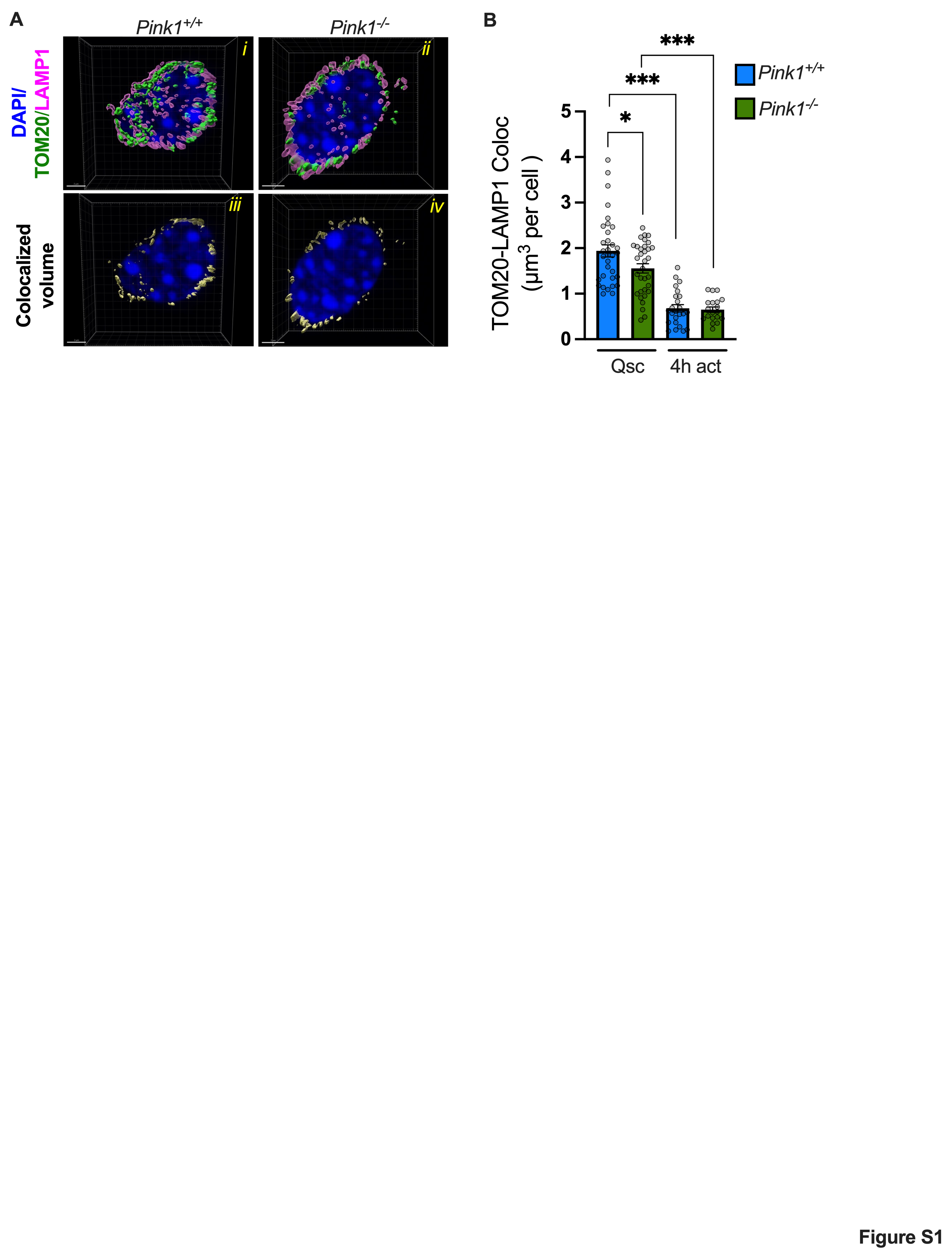
